# Generalised empirical Bayesian methods for discovery of differential data in high-throughput biology

**DOI:** 10.1101/011890

**Authors:** Thomas J Hardcastle

## Abstract

**Motivation:** High-throughput data are now commonplace in biological research. Rapidly changing technologies and application mean that novel methods for detecting differential behaviour that account for a ‘large *P*, small *n*’ setting are required at an increasing rate. The development of such methods is, in general, being done on an *ad hoc* basis, requiring further development cycles and a lack of standardization between analyses.

**Results:** We present here a generalised method for identifying differential behaviour within high-throughput biological data through empirical Bayesian methods. This approach is based on our `baySeq` algorithm for identification of differential expression in RNA-seq data based on a negative binomial distribution, and in paired data based on a beta-binomial distribution. Here we show how the same empirical Bayesian approach can be applied to any parametric distribution, removing the need for lengthy development of novel methods for differently distributed data. Comparisons with existing methods developed to address specific problems in high-throughput biological data show that these generic methods can achieve equivalent or better performance. A number of enhancements to the basic algorithm are also presented to increase flexibility and reduce computational costs.

**Availability:** The methods are implemented in the **R** `baySeq` (v2) package, available on Bioconductor http://www.bioconductor.org/packages/release/bioc/html/baySeq.html.

## 1 INTRODUCTION

High-throughput data are becoming ubiquitous in biological research and numerous statistical techniques have been developed to analyse these data, often to identify patterns of difference between sets of biological replicates. Microarray technology led to a proliferation of methods (Murie *et al.*, 2009) designed to analyse data with many features (i.e., genes) but few biological replicates (the ‘large *P*, small *n*’ problem (Johnstone and Titterington, 2009)) under an assumption of (log-)normality.

The subsequent emergence of count data from high-throughput sequencing (HTS) experiments motivated the development of analysis methods which generally assume some form of over-dispersed Poisson distribution (Soneson and Delorenzi, 2013). The majority of these analytic methods seek not merely to adjust for the high-dimensionality of the data (Benjamini and Hochberg, 1995), but to exploit it through various forms of information ‘borrowing’ across the *P* dimension. However, many of the methods developed achieve this borrowing of information by exploiting specific features of the data. Consequently, while the methods developed for analysis of negative binomially distributed HTS data are conceptually similar to those previously developed for analysis of (log-) normally distributed data, their implementation is very different.

Novel technologies for high-throughput generation of biological data may require different distributional assumptions to current methods. Complete analyses of single-cell sequencing seem likely to require novel distributional assumptions (Brennecke *et al.*, 2013; Islam *et al.*, 2011), as do analyses of high-throughput quantitative proteomic and metabolomic data (Ewald *et al.*, 2009; Nilsson *et al.*, 2010). In addition, complex experimental designs are producing diverse types of data from a single organism (Wang *et al.*, 2014; Yu *et al.*, 2014) which require the development of methods for multi-dimensional and integrative analyses. While some of these challenges are beginning to be addressed, this is being done on an ad hoc basis, requiring further development cycles and a lack of standardisation between analyses.

We present here a generalised method for identifying differential behaviour applicable to high-throughput data of any type. This approach is based on our `baySeq` algorithm, initially developed for identification of differential expression in RNA-seq data using a negative binomial distribution (Hardcastle and Kelly, 2010), and later adapted to paired data using a beta-binomial distribution (Hardcastle and Kelly, 2013). These algorithms extended the method of parametric empirical Bayes point estimation (Morris, 1983) by using multiple point estimators sampled across the ’large *P*’ dimension to approximate a distribution on the hyper-parameters of the data. Here we show that this empirical Bayesian approach (`baySeq v2`) is applicable to any parametric distribution, removing the need for time-consuming development of novel methods for each new type of data.

This generalisation allows differential behaviour to be identified in any class (or combination of classes) of biomolecular event detectable by the application of high-throughput technologies. Here, we define ‘biomolecular event’ to describe a biomolecular process producing a measurable signal with the potential to vary between experimental conditions. This may involve measurement of levels of biomolecules; for example, mRNA-seq or high-throughput proteomics (Nilsson *et al.*, 2010), modifications to biomolecules; for example, methyl-seq and chIP-seq, or levels of interaction between biomolecules; for example, ribo-seq (Ingolia *et al.*, 2009).

To assess the utility of this approach, we consider analyses on microarray data, assumed to be log-normally distributed, and simulated zero-inflated negative binomial data, distributions novel to the empirical Bayesian approach here described. We compare the performance of `baySeq v2` to specific methods developed for analyses of such data in identifying differential expression. Comparisons on over-dispersed Poisson data are largely superfluous since `baySeq` was originally designed for such data, and extensive independent comparisons (Kvam *et al.*, 2012; Cordero *et al.*, 2012; Rapaport *et al.*, 2013) have already been carried out. Nevertheless, to demonstrate that the performance of `baySeq v2` has not been degraded by the generalisation we include a re-analysis of the Rapaport *et al*. (2013) data in Supp. Fig. S1. We additionally demonstrate the capability of `baySeq v2` to perform various novel analyses on a complex set of RNA-seq data from matched tissue sampling in four age groups of rat (Yu *et al.*, 2014). Code examples describing these analyses are included in Supp. S7.

## 2 METHODS

We consider the first dimension of the data to define a specific biomolecular event, and define the data attached to a particular biomolecular event *c* as *D_c_*. Thus, for the simple case of mRNA-seq, the *D_c_* describes the number of sequenced reads for a gene *c* in each biological sample. The second dimension of the data gives an indexing of the samples; thus, *D_cj_* refers to data from the *j*th sample for the *c*th biomolecular event. Further dimensions of the data may be used to refer to individual components of a biomolecular event; e.g., timepoints or marker classification. In addition to the sequenced (or other stochastic) high-throughput data, we may also consider *observables*. These are known, fixed observations that influence the generation of the data. Typical examples of these observables include library scaling factors (a measure of the depth of sequencing for each sample), coding sequence length (in mRNA-seq experiments), and cytosine to uracil non-conversion rates (in bisulphite sequencing data).

As in Hardcastle and Kelly (2010, 2013), we suppose that there exists some model *M* whose posterior likelihood, given the observed data, is to be estimated. The model is defined by the equivalence classes {*E*_1_,…, *E_m_*} such that samples *i* and *j* are equivalent for biomolecular event *c* if and only if *D_ci_* and *D_cj_* are drawn from distributions with identical parameters. For notational simplicity, we define the set 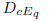 as the data associated with equivalence class *E_q_*. We similarly define the replicate sets {*F*_1_,…, *F_R_*} such that samples *i* and *j* are in the same replicate set *F_r_* if and only if they are biological replicates, and define the set 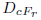 as the data associated with replicate set *F_r_*.

As an example of model construction on these lines, consider an experiment in which three biological conditions *A*, *B* and *C* exist, each with two biological replicates *A*_1_, *A*_2_, *B*_1_, *B*_2_, *C*_1_, *C*_2_. The replicate sets then consist of *F*_1_ = {*A*_1_, *A*_2_}, *F*_2_ = {*B*_1_, *B*_2_}, *F*_3_ = {*C*_1_, *C*_2_}. The set 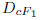 is the observed data of the *c*th biomolecular event in samples *A*_1_ and *A*_2_. Models may be defined encompassing no differential expression, differential expression between all three conditions, and differential expression of one of the biological conditions relative to the other two; for example, the model describing biomolecular events in which condition *A* is different to both *B* and *C*, while *B* and *C* behave equivalently, is defined by equivalence classes *E*_1_ = {*A*_1_, *A*_2_} and *E*_2_ = {*B*_1_, *B*_2_, *C*_1_, *C*_2_}.

The posterior likelihood of a model *M* for biomolecular event *c* is then acquired by computation of

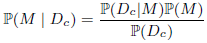

The major challenge in estimating the posterior likelihood of *M* is in estimating ℙ(*D_c_*|*M*), the likelihood of the data for a particular biomolecular event *c* given the model. If *θ_q_* is a random variable defining the parameters of the distribution of the data for the *q*th equivalence class 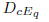 and we assume that the *θ_q_* are independent with respect to *q*, then

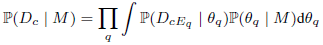

We refer to the joint distribution of *θ_q_* as the hyperdistribution of the data 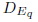. If *Θ_q_* is a set of values sampled from the distribution of *θ_q_*, then the integral in Eqn. 2 can be numerically approximated (Evans and Swartz, 1995) to give

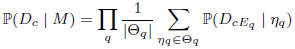

Since we assume that the observed data are independent, given the *η_q_*, this becomes

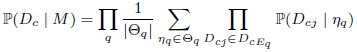

and hence the definition of ℙ(*D_cj_* | *η*), the density function of the data observed for a single sample is sufficient for the posterior likelihoods of the models to be calculated using this framework, provided the sets *Θ_q_* can be constructed.

We further note that posterior distributions on *θ_q_*, given a model *M* and the observed data 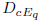, can be estimated by weighting each 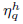 in *Θ_q_* by 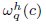, where

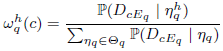

### Construction of *Θ_q_* by sampling from data

Given a density function *f*(*D*; *η*), a model *M*, and a replicate structure {*F*_1_, *F*_2_, ⋯, *F_R_*}, the sets {*Θ*_1_,…, *Θ_m_*} are acquired by sampling from the data. It is often convenient to assume that certain parameters of the distribution of the data are (marginally) identically distributed under all circumstances. In negative binomial modelling of high-throughput sequencing data, for example, the dispersion is commonly assumed to be fixed for any given transcript. This strategy reduces the number of parameters to be estimated from the data and, especially for low numbers of replicates, will tend to increase the stability of the estimated values. We thus categorise the *η_ql_* as either marginally identically distributed over all *q* and models *M*, or not.

We first estimate the marginally identically distributed *η_ql_* using the defined replicate sets. Suppose that we sample the data for some biomolecular event *h*. We first consider the likelihood of the data as the product of the likelihood of the data within each replicate group

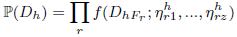
and choose the 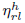 to maximise this likelihood subject to the constraint that 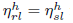 for all *r*, *s* if 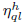 is assumed to be marginally identically distributed over all *q* and models *M*.

For each equivalence class *E_q_*, we then estimate the non-marginally identically distributed 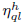 by calculating

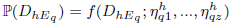
and maximise this likelihood subject to the constraint that 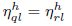 for all *l* if 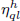 is assumed to be marginally identically distributed over all *q* and *M*. This gives a single sampling of values for each 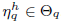. We continue sampling (without replacement) to acquire sufficiently large *Θ_q_*.

In both maximisations, we use the Nelder and Mead (1965) algorithm as implemented in **R**’s `optim` function. This requires initial values to be provided. For optimal performance, these initial values should be as close as possible to the solution to the optimisation, and so `baySeq` v2 allows these to be specified as a function of the sampled data *D_h_*. In practice, maximum likelihood solutions will not always be optimal. In certain circumstances we find increased performance by constraining the domain of the function to be optimised. We give an example of this below when considering a zero-inflated negative binomial model.

If the sampled data *D_h_* derive from a biomolecular event belonging to a model *M*, then parameter estimates from this data for other models may not reflect the true underlying distribution of parameters. For example, data from truly differentially expressed genes will tend to lead to estimates of high variability if it is assumed that they are not differentially expressed. The estimated distribution of parameters for a model can thus be further refined through a weighting of the sampled 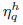 by 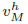, the likelihood that biomolecular event *h* is explained by that model. In this case, Eqn. 4 becomes

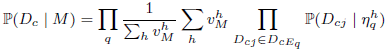

Aquisition of the weights 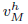 through a bootstrapping strategy is described in Supp. S1. Weighting the sampled 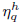 allows comparisons between two models with identical equivalence class definitions but with different hyperdistributions, as described in Supp. S2.

### Model priors

The model priors ℙ(*M*) may be provided based on prior knowledge, or (by default) estimated empirically from the calculated ℙ(*D_c_* | *M*) values. If estimated empirically, the default behaviour of `baySeq` v2 is to calculate the ℙ(*D_c_* | *M*) for all models *M* and all *c* and use the Bayesian Information Criterion (BIC) to choose between each model for each *c*. The proportion of biomolecular events for which a model *M* is selected using the BIC is taken as the prior value ℙ(*M*). If no events are selected for a given model, the prior value for that model is set to 1/*C*, where *C* is the total number of biomolecular events (the prior that would be used if a single biomolecular event were selected to conform to that model via the BIC), and all priors are then scaled to sum to 1. The use of the BIC gives better estimates (Supp. Fig. S2) of the number of differentially expressed biomolecular events than the iterative method described in previous work (Hardcastle and Kelly, 2010).

Rather than assume a single value for ℙ(*M*), `baySeq v2` now allows different subsets of the data to take different values for the model prior. This can substantially improve performance (Supp. S3) if there are strong reasons to suppose that different subsets of the data will display different proportions of differential expression. This may be valuable in a variety of cases where sufficient information is available to distinguish between large categories of genes, for example, if a transcription factor is known to bind to a specific set of gene promoters, this subset of genes is much more likely than its inverse to be differentially expressed if this transcription factor is misregulated. Some care may be needed with this approach in avoiding confirmation bias in downstream analyses of the sets of differentially expressed data.

### Computational Strategies

Calculating priors using numerical methods and posterior likelihoods via Eqn. 3 or Eqn. 8 are computationally expensive steps that scale linearly with both the number of models to be evaluated and the number of biomolecular events being considered. Several strategies are proposed to mitigate the computational costs involved.

#### Stratified Sampling

A minimum size of the sets *Θ_q_* is required for accurate estimation of the posterior likelihoods. The highest accuracy will generally be obtained by making *Θ_q_* as large as possible, but this carries computational costs, making sampling from the data necessary. However, for the numerical approximation described in Eqn. 3 to provide a reasonable approximation to the true value of ℙ(*D_c_*|*M*), the sets *Θ_q_* must contain values in the high probability mass regions of 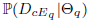. If the sets *Θ_q_* are acquired by sampling uniformly from the data, this can present difficulties for estimating posterior probabilities for 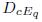 that lie in the tails of the hyperdistribution. Increasing the sample size will resolve this issue, but at a computational cost. Instead, we propose a stratified sampling technique in which the data are stratified by some summary statistic and approximately equal volumes of data are sampled from within each stratum. A sampling 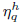 is then weighted by *s^h^*, where *s^h^* is proportional to the size of the stratum from which the value 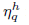 is sampled divided by the number of samplings from that stratum, scaled such that ∑*_h_ s^h^* = 1. Thus, the smaller the proportion of data sampled from a stratum, the greater the weighting applied to the parameters derived from this data. Eqn. 8 then becomes

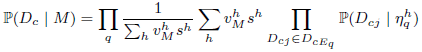

Supp. Fig. S3 illustrates the application of a stratified sampling and shows the effect of performance in simulated data. The primary effect of stratified sampling in these data is to reduce false discoveries for low numbers of selected differentially expressed genes.

#### Consensus Priors

For large numbers of models, computational costs can be reduced substantially if we assume that the parameters are identically distributed for all models; that is, that *Θ_q_* = *Θ* for all *q*. In this case, Eqn. 4 becomes

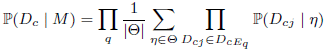
and Eqn. 9 becomes

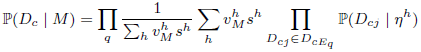

The advantage of this formulation is that the values ℙ(*D_cj_* | *η*) are identical for all models. Consequently, these need be calculated only once to allow the likelihood of the data under any model to be evaluated with the appropriate product-sum-product, considerably reducing the computational cost.

In estimating a set *Θ*, those parameters assumed to be marginally identically distributed over all *q* and models *M* are estimated as previously described in Eqn. 6. We then randomly select amongst the replicate sets a single set *F_r_* and maximise the likelihood

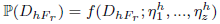
as in Eqn. 7, subject to the constraint that 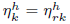 for all *k*, if 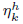 is assumed to be marginally identically distributed over all *q* and *M*. This gives a single sampling of values for each *η* ∈ *Θ*.

## 3 RESULTS

### Affymetrix Microarray Latin Square Data

Microarray data have conventionally been analysed under an assumption of (log) normality. We compare the performance of `limma` (Smyth, 2004), a well established method for discovery of differential expression, to that of `baySeq v2` using a normal distribution in which

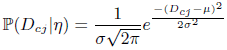

where *η* = (*μ*, *σ*) with the distribution of the standard deviation *σ* being assumed to be constant across samples.

Comparisons are made using the Affymetrix HGU133A Latin Square data (Affymetrix, 2002). These data consist of three technical replicates of 14 hybridisations in a human background, with 42 spiked transcripts at differing concentrations in each hybridisation. We processed the data using the RMA Irizarry *et al.* (2003) algorithm using the alternate chip description file supplied with the data. Non-differentially expressed control spikes showed highly variable expression across arrays (Supp. Fig. S4) and were removed from further analysis.

To assess performance of the methods, we select seven non-overlapping pairs of hybridisations in which to identify pairwise differential expression, and compute the average numbers of true and false positives across these independent pairs of hybridisations. We repeat the selection of pairs of hybridisations one hundred times, and show the distribution of false positives against oligonucleotides selected (Fig. 1a) and of ROC curves (Fig. 1b). The ROC curves are almost identical between the two approaches, while the numbers of false positives in the top *N* transcripts are slightly lower in the `baySeq v2` analysis, suggesting that a generalised empirical Bayesian approach can match or exceed the performance of a well-established method for microarray analysis. False discovery rate (FDR) prediction is considerably better in `baySeq v2`, with the `limma` Benjamini and Hochberg (1995) adjusted p-values predicting no true positives (Supp. Fig. S5).

**Fig. 1.**
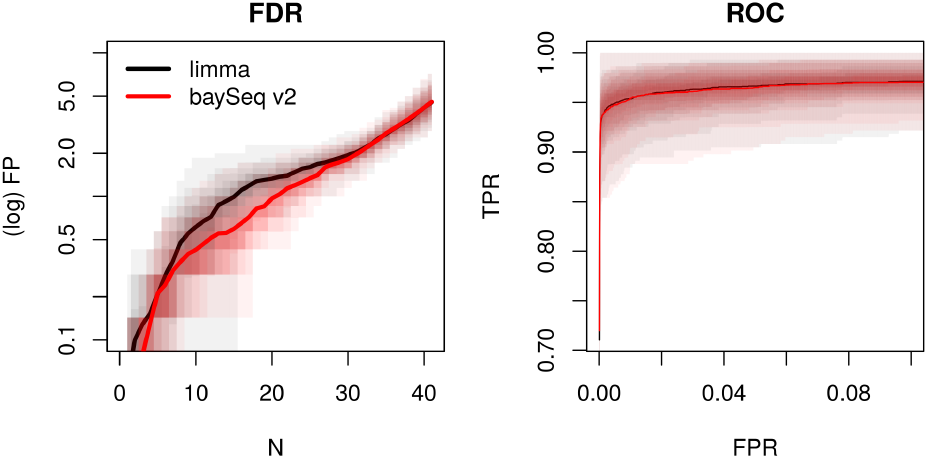
Average numbers of false positives in the top *N* transcripts (a) and ROC curves (b) for `baySeq v2` and `limma` analyses of samplings of non-overlapping pairs of hybridisations in the Affymetrix HGU133A Latin Square data. Percentiles of false discovery rates across samplings are shown as transparent areas around curves.

### Zero-inflated RNA-Seq data

Zero inflation occurs when two processes are involved in the generation of data. The first, a binary distributed process defines whether signal is present or absent, the second generates a distribution on the size of the signal (which may itself be zero) if a signal is present. Zero-inflated negative binomial data may arise in a number of ways in high-throughput sequencing technology. In cross-species analyses (Brawand *et al.*, 2011) in which the expression of gene homologues is being compared, some genes may have moved out of a given regulatory pathway and be non-expressed in some organisms. In meta-transcriptomic studies, (Fang *et al.*, 2014) the observed expression of a gene may be driven by a single organism which may or may not be present in the meta-sample. Similarly, in single-cell sequencing, the expression of genes may be much more of a stochastic on/off process than observed in a multi-cell profile (McDavid *et al.*, 2013). Zero-inflation may also occur in genome-wide enrichment data as a result of low coverage and sequencing bias (Rashid *et al.*, 2011).

The ShrinkBayes package (Van De Wiel *et al.*, 2013), a generalised linear model approach, is the only currently existing method for applying a zero-inflated negative binomial (ZINB) model to high-throughput sequencing data. We compare the results from this package (as implemented in Supp. S7.2.3) to those from a `baySeq v2` analysis using a ZINB model in which

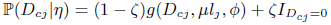

where *g*(*D_cj_*, *μl_j_*, *ϕ*) is the probability mass function of a negative binomial distribution with mean *μl_j_* and dispersion *ϕ*, where *l_j_* is the library scaling factor of library *j*. 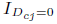 is an indicator function which is 1 if *D_cj_* = 0 and 0 otherwise. *η* = (*μ*, *ϕ*, *ζ*), with the distributions of the dispersion *ϕ* and proportion of zero inflation *ζ* being assumed to be constant across samples. In the event that no zeros appear in the reported expression for a gene, a maximum likelihood estimation of the *ζ* parameter (Eqn. 6) will be zero (up to computational precision). Similarly, since a highly dispersed negative binomial variable will be rich in zeros, a maximum likelihood estimation for a zero-inflated gene may report high *ϕ* and low *ζ* values. Performance is improved (Supp. Fig. S6) by limiting the domain of the likelihood function such that 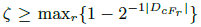, that is, *ζ* must be greater or equal to that proportion of zero-inflation which gives a 50% chance of seeing no zeros within the smallest replicate group. The incorporation of this constraint is shown as a code example in Supp. S7.2.2.

In the absence of a zero-inflated data set for which true positives are known, comparisons are made on simulated data described in Supp. S4. These data are simulated from ten sequencing libraries and contain one thousand ‘genes’ differentially expressed between the first five samples and the second five samples, as well as nine thousand non-differentially expressed genes. We compare the performance of `baySeq v2` with a ZINB model and with a negative-binomial model to ShrinkBayes, implemented with non-parametric priors using default settings. Results from an implementation of ShrinkBayes with mixture-model priors is shown in Supp. Fig. S6. Fig. 2 shows average ROC curves calculated from the simulations, with the mean expression scaled by 1, 3, and 5 in order to explore the effects of increased sequencing depth in a zero-inflation scenario. In all cases, `baySeq v2` with a zero-inflated negative binomial model outperforms ShrinkBayes, especially for low sequencing depths. ShrinkBayes generally outperforms `baySeq v2` with a negative binomial model. There is a general improvement in performance with increased sequencing depth for the two methods accounting for zero inflation which is reversed for the method which does not, suggesting that zero inflation becomes increasingly significant with higher sequencing depths, as might be expected. FDR prediction (Supp. Fig. S7) is considerably better for the zero-inflated model of `baySeq v2`, with strong correspondence between true and predicted FDRs. ShrinkBayes mixture model priors provide better FDR prediction than do the non-parametric priors, but both substantially underestimate the true false discovery rate. Control of FDR was examined by simulating sets of data with no truly differentially expressed genes (Supp. Fig. S8), with all methods showing reasonable control over FDR in this case.

**Fig. 2.**
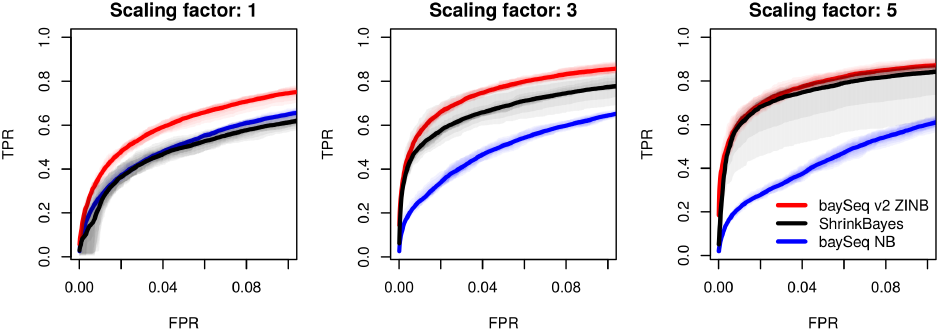
Average ROC curves for `baySeq v2` (ZINB), ShrinkBayes and `baySeq v2` (negative binomial) differential expression analyses in zero-inflated negative-binomially distributed data. Library scaling factors are increased by factors of 1, 3, and 5. Percentiles of true positive rates across samplings are shown as transparent areas around curves.

### Matched sample sequencing

The Rat BodyMap (Yu *et al.*, 2014) project generated RNA-seq data from multiple organs from juvenile, adolescent, adult and aged Fischer 344 male and female rats. For each individual in this study, mRNA is sequenced from every available organ. The expression of any gene across multiple tissues in the same individual is likely to be non-independent since an individual-specific effect may have a global influence on that gene’s expression. We assume that the individual-specific effects may be approximated by some fold-change consistently applied across tissues, and that consequently, the ratio of expression between tissues will remain approximately constant within biological replicates. We thus use these data to demonstrate a novel analysis which allows us to identify changes in relative expression within the tissue types while accounting for individual-specific effects.

Data from ten tissue types (adrenal gland, brain, heart, kidney, liver, lung, muscle, spleen, thymus, and uterus) in female rats are compared between four juvenile (2-week old) to four aged (104-week old) individuals. The data are thus multi-dimensional; for each gene and each individual, we have ten values giving the expression in each organ. A `baySeq v2` analysis is constructed using a Dirichlet-multinomial analysis in which *η* = (*p*_1_, *p*_2_, *ϕ*). The distribution of *ϕ*, the dispersion parameter, is assumed to be constant across samples. The values *p*_1_ and *p*_2_ represent the proportion of expression in the tissues with highest and second highest mean expression in the gene being modelled, with the proportion of expression in the eight remaining tissues being modelled as 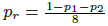, such that

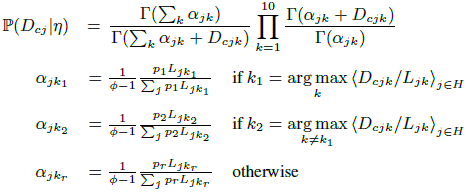

*L_jk_* is the library scaling factor for the *k*th tissue of the *j*th individual and *H* the (model-dependent) subset of samples over which the highest and second highest mean expression tissues are being defined. This reduction in parameters from the full Dirichlet-multinomial is required to control the maximum number of parameters estimated from the eight individuals. Furthermore, this reduces the dimensionality of the distribution being empirically estimated, and hence prevents the empirical distribution from being too sparse an estimate of the true distribution.

We fit three models to these data. The first model describes genes with consistent levels of expression across all tissue types and ages. The second model describes genes with expression consistent between ages, but variable amongst tissue types. The third model describes genes for which the ratio of expression between tissues varies between juvenile and aged individuals. In the first two of these models, all individuals lie in the same equivalence class.

To distinguish between those genes which have consistent levels of expression across all tissue types and ages and those which have consistent levels of expression across ages but vary amongst tissue types, we start by computing priors for a single model of consistent expression between ages. For the model of consistent levels of expression across all tissue types and ages, we take the computed dispersion parameters and set 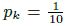 for all *k*, while for the model of consistent expression across ages with variable expression among tissues, we use the maximum likelihood estimates of *p_k_*. We initially weight the models by partitioning the values *p*_1_ estimated for a model of consistent expression between ages (Supp. Fig. S9) to minimise the intra-class variance (Supp. S2) and use Eqn. 8 to calculate posterior likelihoods based on these weighted values. For the model of consistent expression across all tissue types and ages, the initial value for 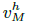 is 1 if the estimated value for *p*_1_ is less than the partitioning threshold, 0 otherwise; for the remaining two models the converse is true. We bootstrap these weightings as described in Supp. S1 over five iterations.

Fig. 3 shows the top ranked genes from each of the three models. The expected number of genes conforming to each model may be inferred by summing the posterior likelihoods estimated for each gene for that model. An estimated 133 genes are expected to be consistently expressed across all tissues and ages. 4603 genes are estimated as showing variability between tissues, but no differential behaviour between ages, while 15430 genes are expected to show variable behaviour in one or more tissues between ages. This unusually high proportion of genes showing differential behaviour is observed since differential expression in any one of the ten tissues will be sufficient to identify differential behaviour.

**Fig. 3.**
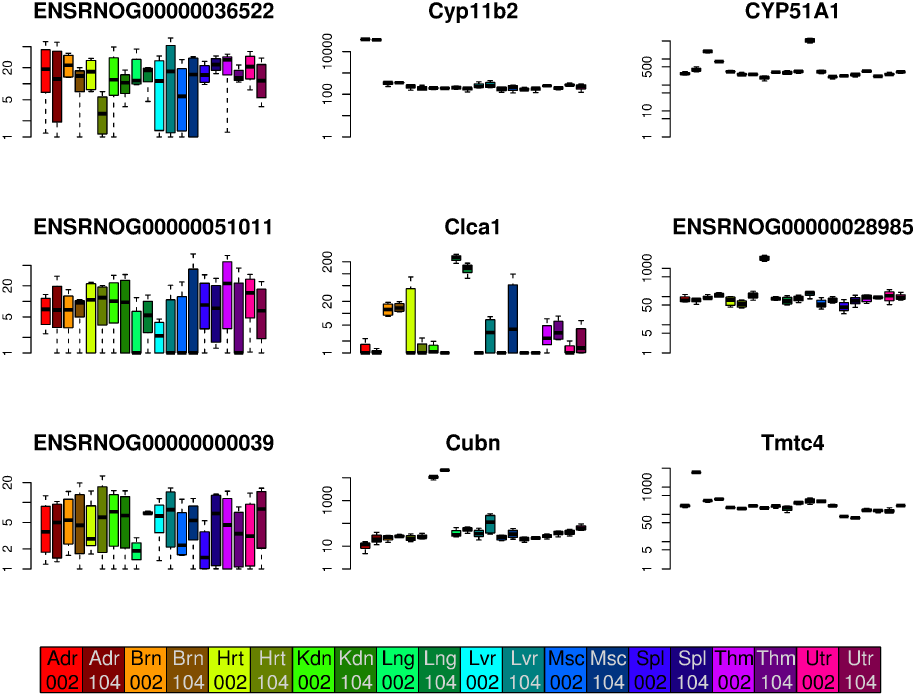
Boxplots of log expression levels of the top three identified genes from female rats for each of three models of expression; consistent expression across tissue types and ages (left), consistent expression across ages but variable between tissues (centre) and variable expression between ages (right). The bar at the bottom of the graphs decodes the colours according to tissue (Adr=adrenal gland, Brn=brain, Hrt-heart, Kdn=kidney, Lng=lung, Liv=liver, Msc=muscle, Spl=spleen, Thm=thymus, Utr=uterus) and age (002=2-week old, 104=104-week old)

Analysis of the estimated posterior distributions of *θ_q_* (Eqn. 5, Supp. Fig. S10) allows a breakdown of the differential behaviour. Of the top 231 genes (selected by controlling family-wise error rate at 10%) that show variability amongst tissues and no differential behaviour between ages, the gene is most abundantly expressed most frequently in brain tissue (36%), and most rarely in uterus tissue (1.2%). Abundances of genes maximally expressed in each tissue correlate well with those reported by Yu *et al.* (2014) as the result of an ANOVA analysis of gene expression (Supp. Fig. S11). Of these 231 genes, 95 show a likelihood greater than 95% of the parameter *p*_2_ exceeding the nominal average proportion of 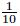. These genes thus show an increase in expression in two tissue types relative to the remaining tissues; the most frequent pairings are between heart and muscle tissues (27) and kidney and liver (15).

For those genes that show a change in proportion of gene expression across tissues over time, we are similarly able to breakdown the discovered differential expression. Controlling family-wise error rate at 10%, we discover 10071 genes that show changes over time. The largest category of change (27%) is a reduction of relative expression in thymus tissue over time, presumably as a result of thymic involution (Shanley *et al.*, 2009). However, in 1073 genes, this reduction in relative expression in thymus tissue correlates with an increase in relative expression in spleen tissue, suggesting a partial compensation mechanism may be in place. The genes showing a reduction in thymus show a strong enrichment for RNA-binding function (Supp. Fig. S12, Supp. Table S1), potentially linked to age-related processes (Masuda *et al.*, 2012). Other large categories of change involve substantial changes in relative expression over time that nevertheless leave the gene maximally expressed in the same tissue (Supp. Fig. S13).

### Complex modelling and computational time

We next use a subset of the Rat BodyMap (Yu *et al.*, 2014) data to demonstrate the use of various computational strategies to carry out a complex modelling analysis. We begin by considering the RNA-seq data for each of the four age groups (2, 6, 21 and 104 weeks) in the thymus of female rats. The total number of potential models for *R* replicate groups is the Bell number *B_R_*. This number scales rapidly with increasing *R*, being bounded above by 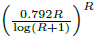 (Berend and Tassa, 2010). For four groups, the total number of models is fifteen, sufficiently low that we are able to evaluate, at some computational cost, posterior likelihoods for each model using Eqn. 8, with the priors being sampled separately for each model as in Section 2. However, we can achieve a significant reduction in computational cost through the use of consensus priors (Eqn. 11).

An alternative way to reduce the computational cost of such analyses is to consider only those models that are biologically interesting, and to employ a ‘catchall’ model to account for data not conforming to one of these models. The catchall model assumes the data for each replicate group is distributed independently, and is thus able to describe any pattern of differential expression reasonably accurately. Consequently, any data not well characterised by any other specified model will thus be best described by the catchall model. Data for which the catchall model has a high posterior likelihood can then be examined for previously unspecified patterns which may be of interest.

We will suppose that we are primarily interested in genes which undergo a single change in expression between two consecutive age groups where this change is maintained in all later age groups. Together with the ‘catchall’, model, and a model for non-differentially expressed genes, this requires the evaluation of five models in total. We refer to these models as NDE (no differential expression), LDE (late differential expression), in which change occurs between the third and fourth age groups, MDE (median differential expression), in which change occurs between the second and third age groups, EDE (early differential expression), in which change occurs between the first and second age groups, and ‘catchall’. We can achieve further reductions in computational cost by using consensus priors in this analysis.

To compare the performance of these approaches, we assume that the estimated posterior likelihoods for the complete fit of the fifteen models without consensus priors are accurate. We can then estimate the number of true positives (and hence, the number of false positives) in each of the models in the restricted analysis for the various approaches as the sum of the posterior likelihoods of the complete fit for the first *N* selected genes. Fig. 4 shows the results of these analyses. There is at most a marginal increase in false discoveries between the complete fit and the complete fit with consensus priors apparent for the NDE set and the EDE set, but in general, the use of consensus priors appears to cause only minor changes in performance for both the complete and reduced model fit. There is a small increase in false positives for the reduced model fit compared to the complete model fit; however, the evaluation of false positives is made under the assumption that the complete model fit is accurate. Consequently, this relatively small difference appears to suggest that both the reduced and the complete model fit are viable alternatives for analysis of specific patterns of expression.

**Fig. 4.**
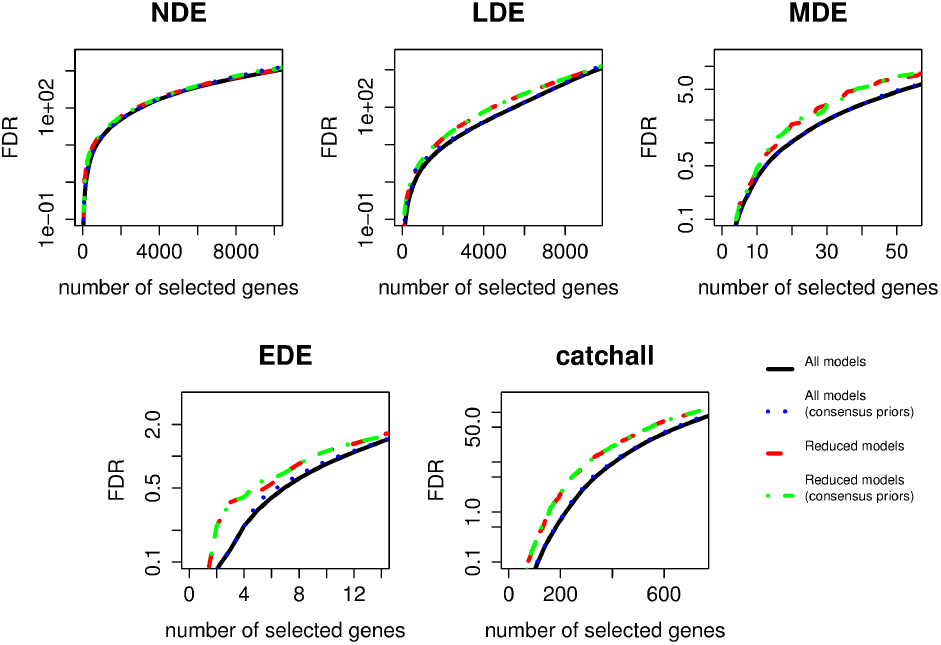
Number of false discoveries estimated in the genes selected by four different strategies for fitting the five models showing conserved change in expression over time.

Using the complete model set permits a greater flexibility in identification of further patterns of differential expression over time. Supp. Fig. S14 shows the expected number of genes for each model, together with normalised and summarised expression values for the top ranked gene in the eight models with highest number of expected genes. These data suggest that, while the majority of genes are not differentially expressed across age groups, almost as many genes show a change between the fourth age group and the three earlier age groups. The models with highest numbers of expected genes generally preserve the ordering of age groups, however, there are exceptional cases for which the second age group is distinct from all other times ({1,3,4}, {2}), and also for which the first and fourth age groups differ from the second and third age groups ({1,4}, {2,3}).

The time required for analysis was evaluated on an octo-core (2.50GHz) machine, running in parallel on all cores. Analysis of the complete model fit took 7.0h, while use of consensus priors reduced this to 1.7h. Analysis of the reduced model took 3.3h, while use of consensus priors in the reduced model took 1.4h. The similarity of performance between the complete model fit with and without consensus priors suggests that consensus priors will generally be preferable for the majority of analyses. It is also apparent from these data that the use of consensus priors scales well, with an increase from 5 models in the reduced model set to 15 in the complete set causing only an 18% increase in computational time.

## 4 CONCLUSIONS

We present here a highly flexible solution (`baySeq v2`) to the general problem of identifying differential behaviour in the ‘large *P*, small *n*’ sets of data that are becoming ubiquitous in biological experimentation (and elsewhere). Given that the parameters of the prior distribution can be inferred from the data, posterior likelihoods for diverse patterns of differential expression can be inferred through an empirical Bayesian analysis. We describe here methods to infer the distribution of the parameters of the prior distribution through maximum likelihood methods, but this is not essential; any method for inferring these parameters from the data might be applicable. In most cases, the inference of these distributions will require some level of biological replication (although this can be avoided by treating all samples as replicates in parameter estimation). The degree of replication required will depend on many factors, including the assumed distribution, the noisiness of the observed data, and the required sensitivity of analyses. A rule of thumb for reasonable parameter estimation (though without any guarantee of biologically meaningful results) may be to include as a minimum *Q* + 1 samples, where *Q* is the largest number of uniquely valued parameters in any model of interest.

We also introduce a number of further refinements to the basic concept. The use of a consensus empirical distribution removes much of the computational cost of these analyses. We demonstrate this in an analysis of complex gene behaviour in a subset of the rat BodyMap data in which the computational time required for a 15 model analysis of 24750 genes in 16 samples is reduced by 75% through the use of consensus priors, with little change in performance. This analysis identifies replicated changes in patterns of differential expression across age groups and shows that diverse types of differential behaviour are present in these data.

Qualitatively distinct data may be distinguished through a weighting or modification of the empirical values representing the hyperdistribution of the data. We show that this technique allows the identification of non-expressed genes in RNA-Seq data and consistent expression over multiple tissues and age groups in matched samples (Fig. 3). Bootstrapping can improve the weightings assigned to the sampled values and further improve performance. A natural extension of this approach would be to use distinct distributions for the different models, and this approach is currently under development. Further extensions to the basic concept are also possible. One of the most advantageous is likely to be the introduction of distributions on parameters estimated across all biomolecular events for a given sample, or within a study. For example, scaling factors involved in RNA-seq analysis are currently treated as observed and fixed values for each library. However, these scaling factors are in fact a random variable whose distribution might be approximated from the observed data. Estimations of this type might be made through bootstrapping or EM approaches, and incorporated into estimates of posterior likelihood.

We demonstrate the effectiveness of the `baySeq v2` approach by comparison with methods designed specifically for particular distributional assumptions. The `limma` (Smyth, 2004) method is a well-established and widely used method for analysis of microarray data assuming a log-normal distribution. We show on the Affymetrix HGU133A Latin Square data a minor improvement in performance of `baySeq v2` over `limma`, under the same distributional assumptions (Fig. 1). In simulated zero-inflated data, `baySeq v2` showed gains in performance over the ShrinkBayes method (Van De Wiel *et al.*, 2013), an approach specifically designed for zero-inflated negative binomial data. These comparisons do not necessarily imply that the accuracy of `baySeq v2` will always match or exceed that of a method specifically designed for a particular set of distributional assumptions, but they do suggest that performance will generally be acceptable.

This approach allows a substantial reduction in development time for information ‘borrowing’ analyses in a ‘large *P*, small *n*’ setting upon data with novel distributional assumptions. This reduction in development time is essential if statistical analysis methods are to keep pace with the rapid development of new technologies and new applications of those technologies that generate large volumes of biological data. For example, the data from single-cell sequencing appears to include a mixture of Bernoulli and Poisson noise (Brennecke *et al.*, 2013), and are likely to require specific distributional assumptions to account for heterogeneity of expression within an individual (Islam *et al.*, 2011). The diverse classes of histone modification signatures (Bernstein *et al.*, 2012) suggests that differential behaviour in histone modification between samples might be identified by the simultaneous analysis of quantitative values for all histone modifications, perhaps through an assumption of a Dirichlet-multinomial distribution. We describe the development of such a model here in the context of analysing multiple matched samples in RNA-seq data from diverse tissues of rat (Yu *et al.*, 2014) in different age groups. The results from this analysis broadly correspond to known interactions between tissues and their changes over time, and allow detailed comparison of gene behaviour between tissues. The relative ease with which distributional assumptions can be changed and modified using these methods also allows the rapid incorporation of significant observables into the models; for example, GC-content, secondary structures or mapping uncertainties.

The second major advantage of this approach is that it allows a standardisation of output across diverse data-types, in that outputs consist of a set of similarly generated posterior likelihoods. Furthermore, posterior likelihoods are easily manipulated and compared between analyses. For example, if a set of RNA-seq data is analysed under an assumption of negative binomially distributed data and a set of ChIP-Seq data under an assumption of a Dirichlet-multinomial distribution, it is straightforward to calculate (under an assumption of independence) joint likelihoods of specific patterns of differential expression of RNA-Seq and ChIP-Seq, and thus, for example, to order the set of overlapping gene/histone modifications by the likelihood that both are differentially expressed. Coupled with the capability of `baySeq v2` to easily evaluate novel datatypes, this suggests that novel data sets can be readily incorporated with existing analyses. Consequently, the methods we present here allow a well-founded statistical framework for the analysis of heterogenous and diverse sets of high-throughput ’omics’ data, a key requirement for future biological research (Gomez-Cabrero *et al.*, 2014).

## ACKNOWLEDGEMENTS

Krystyna A. Kelly reviewed the manuscript and the `baySeq v2` package and made several valuable criticisms.

*Funding*: This work was supported by European Research Council Advanced Investigator Grant ERC-2013-AdG 340642 - TRIBE.

